# Measuring autofluorescence spectral signatures for detecting antibiotic-resistant bacteria using Thermofisher’s Bigfoot spectral flow cytometer

**DOI:** 10.1101/2024.05.13.593944

**Authors:** Sharath Narayana Iyengar, Valery Patsekin, Bartek Rajwa, Euiwon Bae, Brianna Dowden, Kathy Ragheb, J Paul Robinson

## Abstract

Application of flow cytometry to microbiology has been limited due to inadequate availability of bacterial-specific stains, expensive antibody-based fluorophores, ineffective stain cell permeability and challenges in differentiating bacterial cells from cell debris due to their similarity and their small size. In addition, staining cells demands multiple washing steps which limits the sensitivity of detection due to the cell volume that is up to two orders smaller than typical eukaryotic cells. Further, most flow cytometers are not equipped to handle pathogenic organisms. Autofluorescence-based detection of cells can be a useful method for bacterial detection as multiple washing steps can be avoided and it also reduces the time and cost of using stains. Multiple studies have shown that the autofluorescence in bacterial cells are mainly linked to specific proteins, enzymes, or enzyme cofactors such as Flavin Adenine Dinucleotide (FAD) and Nicotinamide Adenine Dinucleotide (NAD) which are involved in bacterial metabolism. In this report, we present a novel method for differentiation between clinically isolated antibiotic resistant and non-resistant bacteria by utilizing their autofluorescence spectral signatures. We utilized a spectral cytometer known as Bigfoot which is equipped with an integrated biosafety cabinet allowing easy handling of pathogenic organisms unlike any other flow cytometers. Bigfoot also has 9 lasers and 54 fluorescence detectors which we utilize to capture the bacterial spectral autofluorescence signatures. As a proof of principle, we initially stressed different types of bacteria (*E*.*coli and Salmonella sp*.) using gentamicin antibiotics by collecting spectral autofluorescence over different time points. The spectral signatures were compared with the non-stressed bacteria. We observed that the stressed bacteria showed an increase in autofluorescence at distinct excitation (Ultraviolet, Violet, and blue color) and emission wavelengths whereas the non-stressed did not. The same experiments were repeated to compare the autofluorescence signatures between Methicillin-resistant and methicillin susceptible *staphylococcus aureus* (MRSA and MSSA) which were stressed with oxacillin antibiotics. MSSA showed an increase in autofluorescence between 4 – 6 h after exposure to oxacillin. MRSA on the other hand showed no increase in autofluorescence and the autofluorescence between stressed and non-stressed MRSA had similar signatures. This demonstrated that the antibiotic resistant or susceptible strain can be detected by observing the change in autofluorescence signature at specific wavelengths in a few hours. This label-free, quantitative, and resistant-specific autofluorescence spectral signatures from the Bigfoot spectral flow cytometer could potentially be utilized for rapid detection of antibioticresistant strain.

## Materials and methods

### Preparation of bacteria samples

Antibiotic susceptible bacterial strains: *Escherichia coli* (ATCC 25922), methicillin-susceptible *Staphylococcus aureus* ATCC 23235, and *Salmonella enteritis* (ATCC 13076), were used in these experiments. Methicillin-resistant *Staphylococcus aureus* PU54 REAR-16 (human clinical isolate), which is resistant to oxacillin sodium salt 95% (ThermoFisher Scientific): MIC ≥ 4 ug/mL (CDC 2019b), was used . To obtain pure bacterial colonies for autofluorescence experiments, all bacteria were grown overnight at 37°C in in a Fisher Scientific Isotemp incubator on Tryptic Soya Agar (TSA) plates. From the overnight culture, colonies from each organism were inoculated into a final volume of 5 mL of Tryptic Soya Broth (TSB), and the concentration was adjusted to 10^6^ CFU/mL by measuring the initial optical density (OD) using a spectrophotometer, WPA Biowave CO8000 Cell Density Meter to obtain pure bacterial cultures of known concentration.

### Sample preparation for autofluorescence experiments

Specific concentrations of antibiotics: gentamicin (Sigma-aldrich) and oxacillin were prepared by diluting the starting stock concentration of 5 mg/mL using TSB broth. Using stock concentration three different concentrations of gentamicin: 0.25, 0.5, 1 ug/mL and three oxacillin concentrations 2, 4 and 8 ug/mL were prepared. For autofluorescence experiments, a total of 2 mL of working samples were prepared which contained 1mL of specific antibiotic concentration, 0.5 mL of pure bacterial culture tube (refer preparation of bacterial sample solution) and 0.5 mL of pure TSB in each tube. For positive controls, 0.5 mL of pure bacterial culture tube were mixed with 1.5 mL of pure TSB and as a negative control 1mL of antibiotics mixed with 1 mL of TSB broth was used. In the final concentration of antibiotics in the working sample were 0.125, 0.25, 0.5 μg/mL of gentamicin and 1, 2 and 4 μg/mL oxacillin respectively. These working samples were incubated at 37°C incubator and the autofluorescence of bacteria was measured at different time points 0, 2, 4, 6 and 24 h using the Bigfoot spectral flow cytometer.

### Measuring bacterial autofluorescence using Bigfoot

For measuring bacterial autofluorescence, 100 μL of the working sample (refer: Sample preparation for autofluorescence experiments) was added to 900 μL of PBS and introduced into the Bigfoot. Bigfoot settings were set to spectral mode to capture the autofluorescence intensity using 7 lasers (excitation wavelengths: 349, 405, 445, 488-532, 561, 640, and 785 nm) and 54 fluorescence detectors. Spectral data captured at different time points was further analyzed using FCS express 7.0 software.

### Waste and contamination management

Managing and proper disposal of bacterial waste, especially pathogenic or antibiotic resistant organisms is very crucial to maintain an uncontaminated environment. Post bacterial experiment the following washing steps are performed for 5 min each, to sterilize the Bigfoot flow cytometer: Step 1: 10% bleach (Clorox), Step 2. Coulter Clenx (Beckman-Coulter), Step 3: Cavicid (Fisher Scientific), Step 4: Hellmanex solution, Step 5: 70% ethanol. After performing all the above washing steps, clean milli-Q water is processed through Bigfoot and the density plot is analyzed on the Bigfoot. If there are any particles observed, the cleaning steps are repeated until the particles are removed and to ensure sterility. When the bigfoot shuts down, the cytometry undergoes washing steps which further sterilize the system. After the bigfoot shutdown, the waste chamber is separated and exposed to 10% bleach overnight or minimum of 4hr before discarding.

## Introduction

### Antibiotic resistance

Improper or non-judicious use of antibiotics has led to the increase in bacterial resistance globally. This has led to the evolution of superbugs, making the treatment of infection using the majority of antibiotics obsolete. Starting from the development of penicillin resistant *staphylococcus* in the year 1940, resistance to wide range of antibiotics which are commonly referred to as antibiotic resistant (AMR) strains has been reported to methicillin resistance staphylococcus aureus (MRSA), erythromycin streptococcus, gentamicin resistant enterococcus, carbapenem resistant *Enterobacteriaceae*, cephalosporin *staphylococcus* (Ventola 2015; WHO 2021) and others. Among them, MRSA is considered as a serious antibiotic resistance threat by centers of disease control and prevention (CDC) AR report in 2019 (CDC 2019a). MRSA infection can be acquired through hospital associated (HA-MRSA) and community associated (CA-MRSA) (Nandhini et al. 2022). It has been estimated that cases of MRSA infections worldwide range from 7% in HIV infected patient to 70% in community associated MRSA (Ko and Moon 2018; Sabbagh et al. 2019; Siddiqui and Koirala 2023; Ventola 2015). Infection from MRSA has high mortality rate, with increased hospitality stay and costs (Lakhundi and Zhang 2018; Whitby et al. 2001; Wolk et al. 2009). The development of methicillin resistance in *staph aureus* is mainly due to the mutation in penicillin binding protein (PBP2a) produced by the *mecA* gene (CDC 2019b; Hiramatsu et al. 1992; Lakhundi and Zhang 2018). Thus, rapid, and effective diagnosis of MRSA infection is crucial for better treatment and to prevent overuse or misuse of antibiotics which can save lives in the hospitals, help in decreasing the related costs and prevent the development of AMR strains.

### Methods for detecting MRSA

According to CDC, the recommended methods for diagnosing MRSA infection are any of the following methods: 1. broth dilution 2. agar plating 3. disk diffusion or other FDA approved molecular diagnosis tests such as polymerase chain reaction (PCR) (CDC 2019b). For antibiotic susceptibility testing (AST) on *staphylococcus* strains oxacillin and cefoxitin are the two common antibiotics used (CDC 2019b). The present golden standard methods such as agar plating and broth dilution methods are time consuming (72 h) and requires multiple culturing steps (Abdou Mohamed et al. 2021; Baltekin et al. 2017; Gosiewski et al. 2014; Iyengar et al. 2021; Narayana Iyengar et al. 2021a; Vasala et al. 2020). For genomic analysis, such as PCR specific primers are needed and demands the clinicians to narrow down the list of microbes to be tested in the sample (Liu et al. 2019; Walker et al. 2016). Despite the fact that PCR has capabilities of detecting MRSA such as by using *mecA* PCR tests, the information about the bacterial viability cannot be obtained using PCR as DNA from live and dead bacteria will be amplified (Cangelosi and Meschke 2014; Dietvorst et al. 2022; Narayana Iyengar et al. 2021a; Sinha et al. 2018). In addition, *mecA* PCR tests cannot detect border line oxacillin resistant MRSA (CDC 2019b).

### Conventional vs spectral flow cytometer

Conventional polychromatic flow cytometers and sorters are an interesting alternative method for detecting bacteria in the sample as they can perform high throughput analysis with high efficiency sorting (Nolan 2022). Flow cytometers can detect and sort bacteria at a single cell level, live and dead cells can be sorted by using staining and as this technique is non-destructive, viable cells can be recovered for further downstream analysis (Robinson et al. 2023; Shapiro 2003). While performing bacterial analysis using flow cytometers, forward scatter (FSC) and the side scatter (SSC) plots are obtained by usually adding a cell staining step (such as live dead staining) to analyze cells. The FSC vs SSC scatter plots provides information somewhat related to cell size and cell shape/granularity. A gate region on the scatter plot was selected which can be used for sorting cells. Despite all the advantages of conventional flow cytometry to the field of microbiology, there are a number of important limitations. For instance, due to the small size of microbes, it is often challenging to differentiate between bacteria and cell debris (Marcos-Fernandez et al. 2022). In addition, there are other limitations of non-specific binding of fluorescent dyes, lack of availability of specific antibodies and high cost of antibody-based fluorophores (Iyengar and Robinson 2024; Koch and Muller 2018; Marcos-Fernandez et al. 2022; Marutescu 2023; Robinson et al. 2023), Cell detection by measuring fluorescence signals (from staining) are limited by the number of lasers and detectors available in some cytometers (Robinson 2019; Robinson et al. 2005).

In conventional flow cytometers, the signals from the stained cells are captured from their highest emission peaks. However, many fluorescent stains have broad emission signal, and many stains can have similar high emission peaks that lead to signal overlap and limits multiplexing capabilities (Robinson et al. 2023; Robinson et al. 2005). Compensation method of data processing can be performed to subtract the spillover signal from the detector of interest, but this can be very challenging in cases where the signal mostly overlaps or bacteria and media have significant autofluorescence (Loken et al. 1977).

To address these limitations from conventional flow cytometers, spectral flow cytometers has gained more interests mainly because of their multiplexing capabilities. Spectral flow cytometers are equipped with multiple lasers and detectors which collect the entire emission spectra rather than a single band (Futamura et al. 2015; Nolan and Condello 2013). For example, if the sample is stained with multiple fluorophores which have similar overlapping highest emission peak signal but differ in off-peak signals, they can be easily differentiated (Nolan 2022; Novo 2022; Robinson 2019). Instead of compensation in conventional flow, the recorded spectral signatures are unmixed where the undesired signal is subtracted from the target stain and the whole spectra is reconstructed for each parameter (Nolan and Condello 2013; Ortolani 2022; Robinson et al. 2005). The difference between polychromatic and spectral flow cytometry is shown in Fig.1. Thus, by using spectral flow cytometry, multiplex analysis can be performed along with improved data resolution and accuracy.

**Figure 1.**
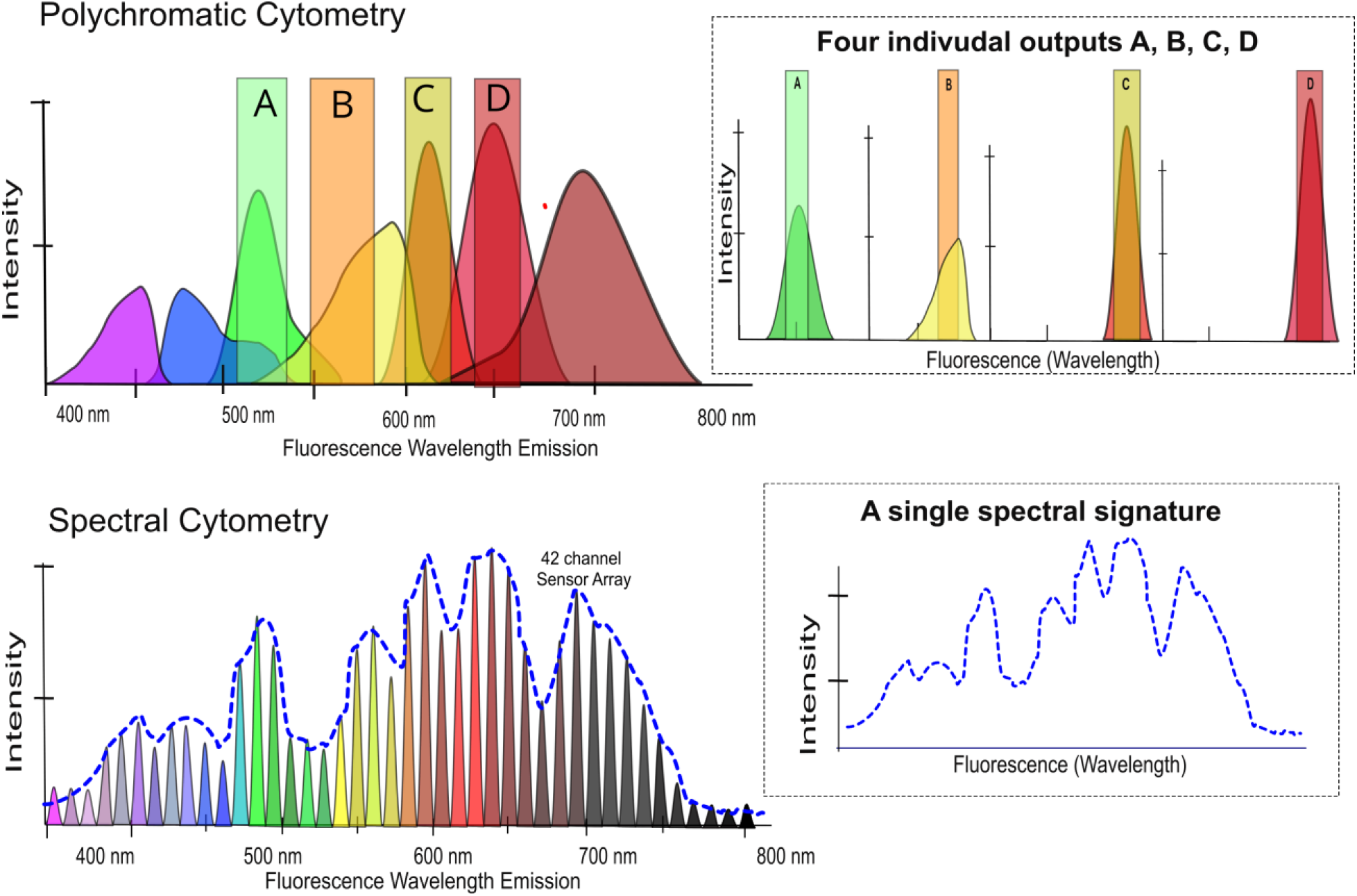
Difference between conventional and spectral flow cytometry. In polychromatic flow cytometry, extracting the actual signal from stains with similar or overlapping peak signals cannot be easily extracted even after performing compensation. On the other hand, spectral flow cytometry can be utilized to obtain the spectral signature across the whole wavelength. By comparing the spectral signature stains with similar overlapping peak signal can be differentiated if they have different off-speak signals.

### Autofluorescence in bacteria

In recent times, detecting and classifying cells based on autofluorescence has gained more interest, especially for immune cells such as T cell subtypes (Walsh et al. 2021). Similarly, detecting bacteria by utilizing their natural autofluorescence signal can provide useful information, as some of the limitations from using fluorescent dyes such as low permeability and high costs of stains can be addressed. The autofluorescence signal from bacteria arises mainly due to the intracellular components such as amino acids (protein structure), lipofuscin, collagen, co-enzymes such as Flavin Adenine Dinucleotide (FAD) and Nicotinamide Adenine Dinucleotide (NAD) which are involved in cell metabolism (Croce and Bottiroli 2014; Surre et al. 2018; Yang et al. 2012). For instance, a study by Mihalcescu et al, showed that by using conventional flow cytometer, the majority of the green fluorescence from *E*.*coli* was mainly linked to flavins: Riboflavin, FAD, FMN and other flavin types (Mihalcescu et al. 2015). Depending on the excitation wavelength, the emission wavelengths of flavins, NAD and lipofuscin emitted in the green, blue, and orange bands respectively (Surre et al. 2018). The peak excitation and emission wavelengths for NADH is between 330 - 360 nm and 360 - 465 nm respectively. For FAD, the excitation and emission wavelengths fall between 440 - 470 nm and 520 - 530 nm respectively (Blacker and Duchen 2016; Blinova et al. 2005; Croce and Bottiroli 2014; Kolenc and Quinn 2019; Skala et al. 2007). Multiple studies have reported that the autofluorescence of bacteria increases when stressed. For example, studies performed by Renggli et al, showed an increase in autofluorescence in *E*.*coli* when exposed to antibiotics such as norfloxacin and ampicillin (Renggli et al. 2013). They also reported how fluorescence-based analysis using conventional flow cytometers can be affected by bacterial autofluorescence. Furthermore, an increase in autofluorescence of *E*.*coli* when stressed with ampicillin was demonstrated by Surre et al, and this increase in autofluorescence was mainly linked to flavins (Surre et al. 2018). It is important to note that all the previous studies observed 1) changes in autofluorescence intensity of stressed bacteria at specific wavelengths using conventional flow cytometers which provide very limited information. 2) all these studies have worked on measuring autofluorescence mainly from *E*.*coli* and other lab-controlled strains. 3) Clinical isolated pathogens have not been explored mainly due to the inability of flow cytometers to handle pathogenic organisms. In this study we demonstrate that we can identify clinically isolated human antibiotic resistant strain by measuring the autofluorescence signature using Bigfoot spectral flow cytometer.

### Bigfoot spectral flow cytometer

ThermoFisher Scientific’s (Waltham, MA) bigfoot spectral flow cytometer is an interesting candidate to study clinical pathogens as they are equipped with extreme biosafety features and has multiplexing capabilities (ThermoFisher 2020).The image of the Bigfoot placed in the normal laboratory settings is shown in Fig.2A. Bigfoot is equipped with 9 lasers and 54 fluorescence detectors to capture spectral signatures and real time spectral unmixing can be performed (ThermoFisher 2020). It also has a separate closed chamber for sheath fluid and waste tank to avoid contamination. Most importantly, bigfoot has a biosafety cabinet similar to biosafety level 2 (BSL-2) with HEPA filters for managing bio contaminant, aerosol (ThermoFisher 2020) as shown in Fig.2B zoomed area. The sample chamber is separate, and six samples can be processed at a single time which allows less interaction of samples and the user. Bigfoot is also a cell sorter which has a separate enclosed sorting chamber for collecting sorted cells. Bigfoot can process sample at very high throughput up to 70,000 events/sec (ThermoFisher 2020). All these features allow safe handling of dangerous organisms such as antibiotic resistant bacteria. An example of the scatter plot (SSC vs FSC) screenshot from Bigfoot for sample containing *E*.*coli* in Tryptic Soya Broth (TSB) media is shown in Fig.2C. A gate is drawn on the scatter plot to select the region of interest and the corresponding spectral signature is captured across the whole spectra. Bigfoot screenshot of the *E*.*coli* autofluorescence spectral signature is shown in the graph (intensity vs wavelength) of Fig.2D. The wavelength here represents the emission wavelength that span from 349 to 785 nm.

**Figure 2.**
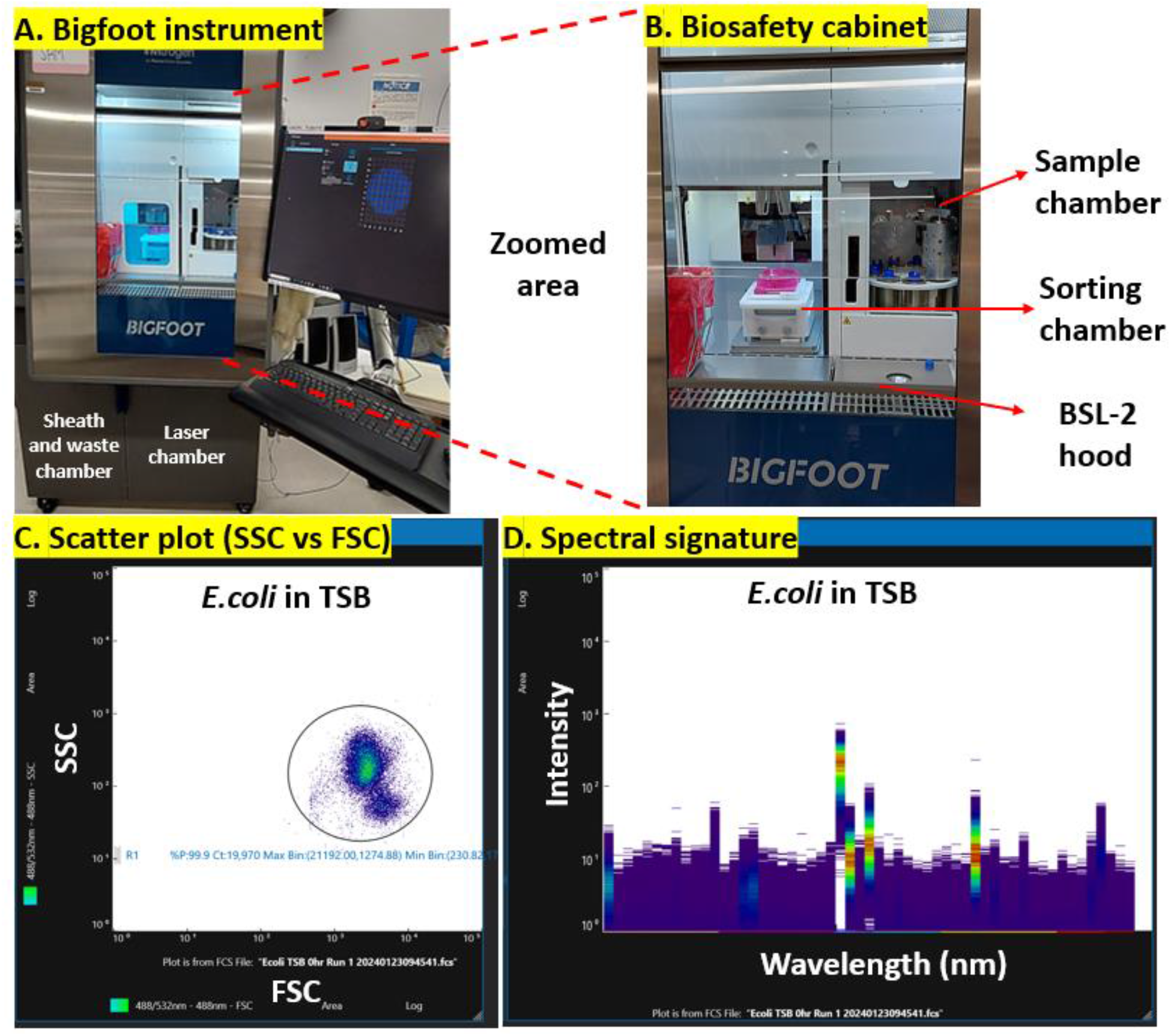
Features of Bigfoot spectral flow cytometer and the process involved in spectral analysis. (A) Image showing the Bigfoot spectral flow cytometer kept in the normal laboratory settings. (B) The zoomed area of the biosafety cabinet which is similar to a BSL-2 hood that allows safe handling of pathogenic organisms is shown. Bigfoot has a separate sample chamber and sorting chamber to ensure user safety and to prevent contamination. (C) Screen shot of the scatter plot from sample containing E.coli in TSB with a gated region for spectral analysis. (D) Screenshot of the autofluorescence spectral signature graph (Intensity vs emission wavelength) of sample containing E.coli in TSB.

## Data analysis

Increased complexity of datasets in many disciplines has demanded data analysis methods that can interpret and reduce large complex data set to smaller dimensions by preserving the original data (Jolliffe and Cadima 2016). Principal component analysis (PCA) is a commonly used method in the research field to analyze and represent multivariable data in an interpretable format (Jolliffe and Cadima 2016; Lever et al. 2017; Migenda et al. 2024). Briefly, PCA is a statistical method where high-dimensional complex data are transformed into a lower-dimensional space defined by principal components (PCs) to account for significant portion of the variance (Greenacre et al. 2022). The new variables (PCs) are the linear functions of the original data points with lower number PCs capturing the highest variance and are uncorrelated to each other (Jolliffe and Cadima 2016). The higher number and the increasing PCs mainly reflect the residual noise (Yeung and Ruzzo 2001). In PCA analysis, the first principal component (PC1) is deduced by projecting the data with the highest variance and thus decreasing the distance between each original data points (Lever et al. 2017). The second principal component (PC2) is orthogonal to PC1 and captures the second highest variance. The succeeding PCs are geometrically orthogonal to all prior PCs and the variance progressively decreases (Lever et al. 2017). Eigen vectors describe the direction of the PCs with maximum variance. Eigen values account for the magnitude of the variance by each PC (Jolliffe and Cadima 2016). Data points with similar properties cluster together on a PCA plot which can be utilized for better understanding of data grouping (Lever et al. 2017; Yeung and Ruzzo 2001). An elaborate description of the PCA method can be found in various literatures (Greenacre et al. 2022; Jolliffe and Cadima 2016; Lever et al. 2017; Migenda et al. 2024).In this work we utilize PCA analysis method to better understand and differentiate between the autofluorescence signatures captured from antibiotic stress and non-stressed bacteria.

The steps involved in detecting antibiotic resistant strain from samples is described in Fig.3. The sample with suspected bacterial infection is inoculated into liquid media (such as Phosphate buffer Saline-PBS) and treated with known concentrations of specific antibiotics. This sample mixture is incubated at 37°C for 4 -6 h. The incubated sample is introduced into Bigfoot spectral flow cytometer. The spectral signatures of the antibiotic stressed bacteria are captured and compared with the non-stressed bacteria (positive control). If the bacteria in the sample mixture shows an increase in autofluorescence compared to the positive control, it suggests that the bacterial strain is non-resistant to the exposed antibiotics. No increase in autofluorescence signal shows the presence of resistant strain in the sample.

**Figure 3.**
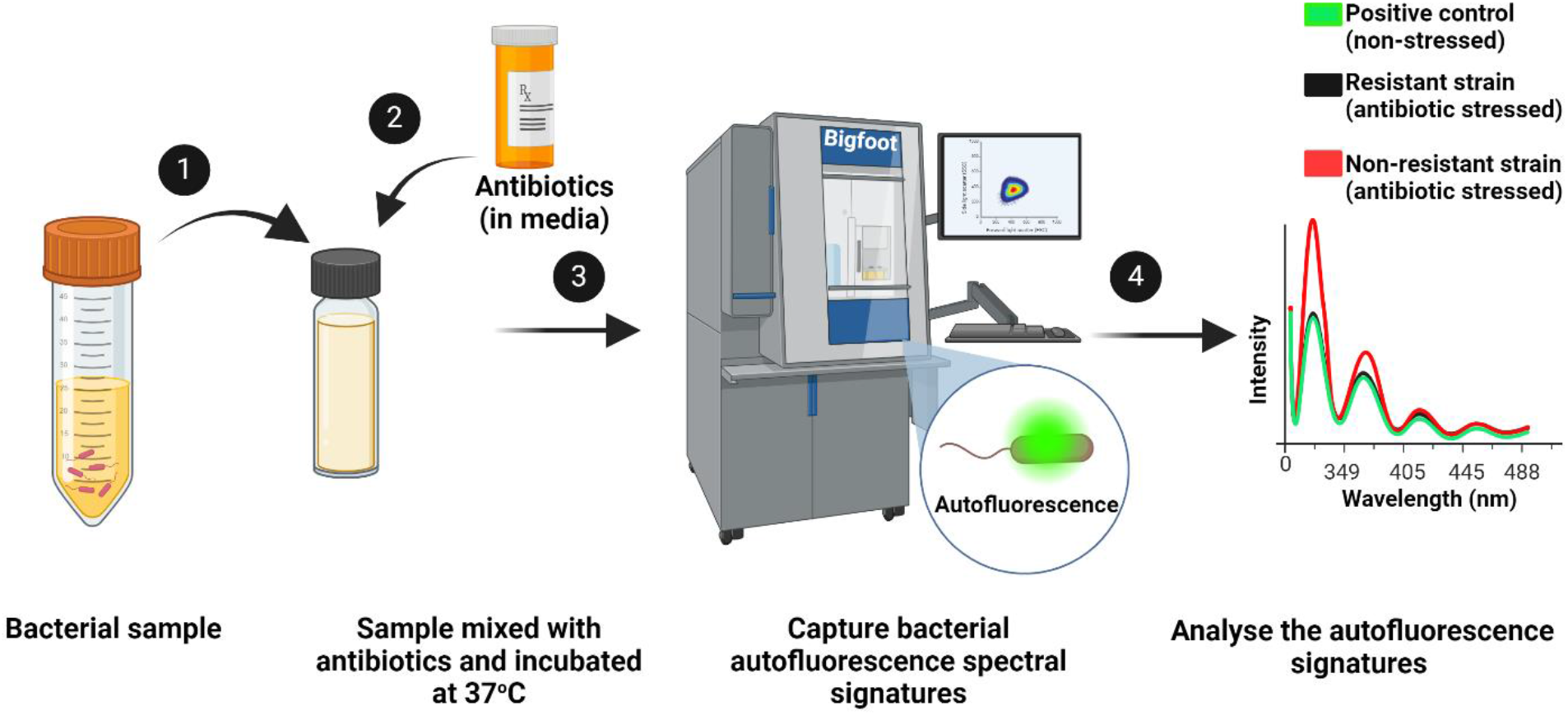
Schematic showing the steps involved in detecting resistant bacteria by measuring bacterial autofluorescence signatures using Bigfoot. Sample containing speculated resistant strain is exposed to antibiotics and introduced into Bigfoot spectral flow cytometer. The autofluorescence spectral signatures is captured and analyzed after 4-6 h. The autofluorescence signal from stressed and non-stressed bacteria is compared. Increase in autofluorescence shows the presence of non-resistant strain while the antibiotic resistant strain shows no increase in the autofluorescence signal.

## Results

### Bacteria stressed with gentamicin antibiotics

As a proof of principle, the effect of gentamicin on bacterial autofluorescence was studied. Two different bacterial species *E*.*coli* ATCC 25922 and *Salmonella enteritidis* ATCC 13076 (SE) were exposed to three different concentrations of gentamicin(0.125, 0.25 and 0.5 ug/mL) where both *E*.*coli* and SE are gentamicin susceptible strains. According to CLSI and CDC, the MIC for *E*.*coli and Salmonella* spp. are in the range of 0.25 – 1 μg/mL respectively (EUCAST 2022). Bacteria grown in TSB media (not exposed to gentamicin) was used as positive controls. *‘TSB media’* and *‘TSB mixed with gentamicin’* was used as negative controls. All the above tubes were incubated at 37°C for 24 h, and the spectral signatures were captured using Bigfoot. The complete protocol is described in the Materials and methods. For each of the tubes, SSC vs FSC plots were captured for both *E*.*coli* and SE as shown in Fig.4A (graphs obtained using FCS express 7.0). Fig.4B shows the overlapped (positive control and sample) unmixed autofluorescence spectral signatures for stressed and non-stressed bacteria obtained using FCS express 7.0 software. Unmixing removes the background spectral signal from negative control: *‘TSB mixed with gentamicin*’. The X axis in the full spectrum graph shows the excitation (Ex) and emission (Em) wavelength combination in the format Ex-Em. For instance, 349 – 387/11 means 349 nm excitation wavelength and 387 center emission wavelengths with 11 nm bandwidth. It was observed that at three specific excitation wavelengths (ultra-violet, violet and blue color) and corresponding emission wavelengths, bacteria stressed with higher concentration of gentamicin (0.5 ug/mL in green) showed increase in autofluorescence (numbered and highlighted with dashed red lines). Increase in autofluorescence were observed with increasing concentration of gentamicin at these specific Ex-Em wavelengths. The graph showing the autofluorescence signature at selected Ex-Em wavelengths (3 lasers - 29 detectors) obtained from the whole spectra is shown in Fig.4C.

**Figure 4.**
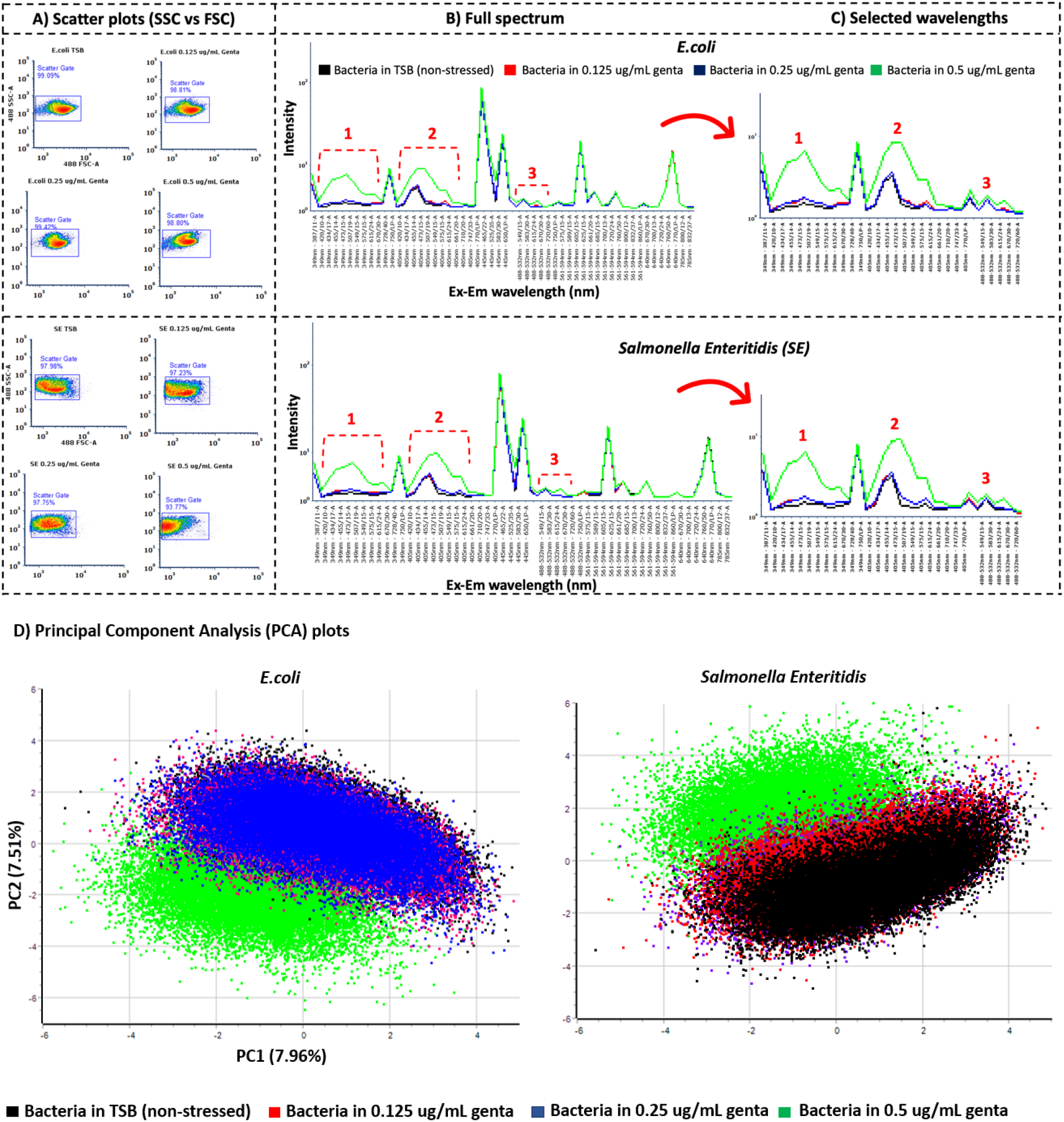
Effect of gentamicin stress on E.coli and Salmonella Enteritidis (SE) autofluorescence recorded using Bigfoot. (A) Side vs forward scatter plots for E.coli and SE exposed to different concentration of gentamicin (0, 0.125, 0.25 and 0.5 ug/mL). (B) Unmixed spectral autofluorescence signature for both types of bacteria analyzed using FCS express. Increase in autofluorescence intensities at three excitations (dashed red lines with numbering) and corresponding emission wavelengths for higher concentration of gentamicin (0.5 ug/mL-green) was observed. (C) Selected wavelengths (3 lasers and 29 detectors) where increase in bacterial autofluorescence were observed from stressed bacteria. (D) PCA plots for E.coli and SE for both stressed and non-stressed case. Shift in PCA clusters for bacteria stressed with high concentration of gentamicin was observed (green – 0.5 ug/mL).

The spectral data at selected wavelengths were further processed using in-house built Cytospec 11.14 software and principal component analysis (PCA) was performed. PCA plots are obtained by projecting the autofluorescence spectral signature obtained from selected wavelengths (3 excitation and 29 emission wavelengths) and projecting them onto two principal components (PC1 and PC2) on a 2D plot based on highest variance. Data points with similar properties cluster together on the PCA plots. Fig.4D shows the PCA plots which is represented in hyper spherical co-ordinate showing the comparison between gentamicin stressed and non-stressed cases for both *E*.*coli* and SE. Each data point on the PCA plot represents the autofluorescence intensity spectral data obtained from 20,000 bacterial cells for both stressed and non-stressed cases. Cells with similar autofluorescence characters cluster together. It was observed that the PCA clusters stressed bacteria showed a clear shift with increasing concentration of gentamicin. A major shift was observed for *E*.*coli* and SE exposed to the highest concentration of gentamicin (green – 0.5 ug/mL).

### Methicillin resistant and non-resistant *staphylococcus* exposed to oxacillin

To study the autofluorescence signatures of antibiotic resistant and non-resistant bacterial strain, experiments were repeated using methicillin suspectable *staphylococcus aureus* ATCC 23235 and clinically isolated methicillin resistant *staphylococcus aureus* REAR 16 strain (MSSA and MRSA). Three separate experiments were performed but one example of the results is shown in Fig.5. Both MSSA and MRSA were exposed to two different oxacillin antibiotic concentrations: 1 and 2 ug/mL, and the autofluorescence spectral signatures were compared. Unstressed MSSA and MRSA (grown in TSB without oxacillin) were used as positive controls whereas TSB containing 2 ug/mL oxacillin was used as negative control. All the samples were incubated at 37°C. In order to deduce the incubation time at which the autofluorescence change, we measured for all the above cases at four different time points 2, 4, 6 and 24 h. The incubation time also represents different growth phase of bacteria. Cytospec 11.14 software was used for data representation and analysis. Fig.5A shows the scatter plots obtained from non-stressed and stressed MSSA and MRSA at 2 and 6 h time points. The scatter plots for 4 and 24 h are shown in supplementary Fig.S2. Changes observed in scatter plots can represent different characteristics of cells such as cell size, cell granularity, cell growth phase etc. It can be observed from the scatter plot that MSSA stressed with oxacillin shows dying cells with increase in the oxacillin concentration from 1 to 2 ug/mL at 2 h compared to MSSA in TSB (non-stressed). This is observed as more cells are found to concentrate closer the origin (0,0) in the graph. Decrease in cell density and cells accumulating at the origin of the scatter plot are mostly due to dead cells or cell debris. Similar observation was found with majority of cells dying for stressed MSSA after 6 h of incubation and with increasing concentration of oxacillin. It is important to note that cells also die naturally as they follow a different growth phase (lag, log and dead phase) as the time of incubation increases. On the other hand, MRSA scatter plots shows no major increase in dead cells exposed to higher concentration of oxacillin (up to 2 ug/mL). Scatter plots can be used as a first step of confirmation to differentiate between MSSA and MRSA sample when stressed with Oxacillin. The scatter plot captures the initial information about the cell behavior in the sample solution when stressed with antibiotics, but it cannot be solely used as a way to differentiate between MSSA and MRSA. The spectral signatures (intensity vs wavelength) comparing the effect of oxacillin on stressed and non-stressed MSSA and MRSA is shown in supplementary Fig.S4.

**Figure 5.**
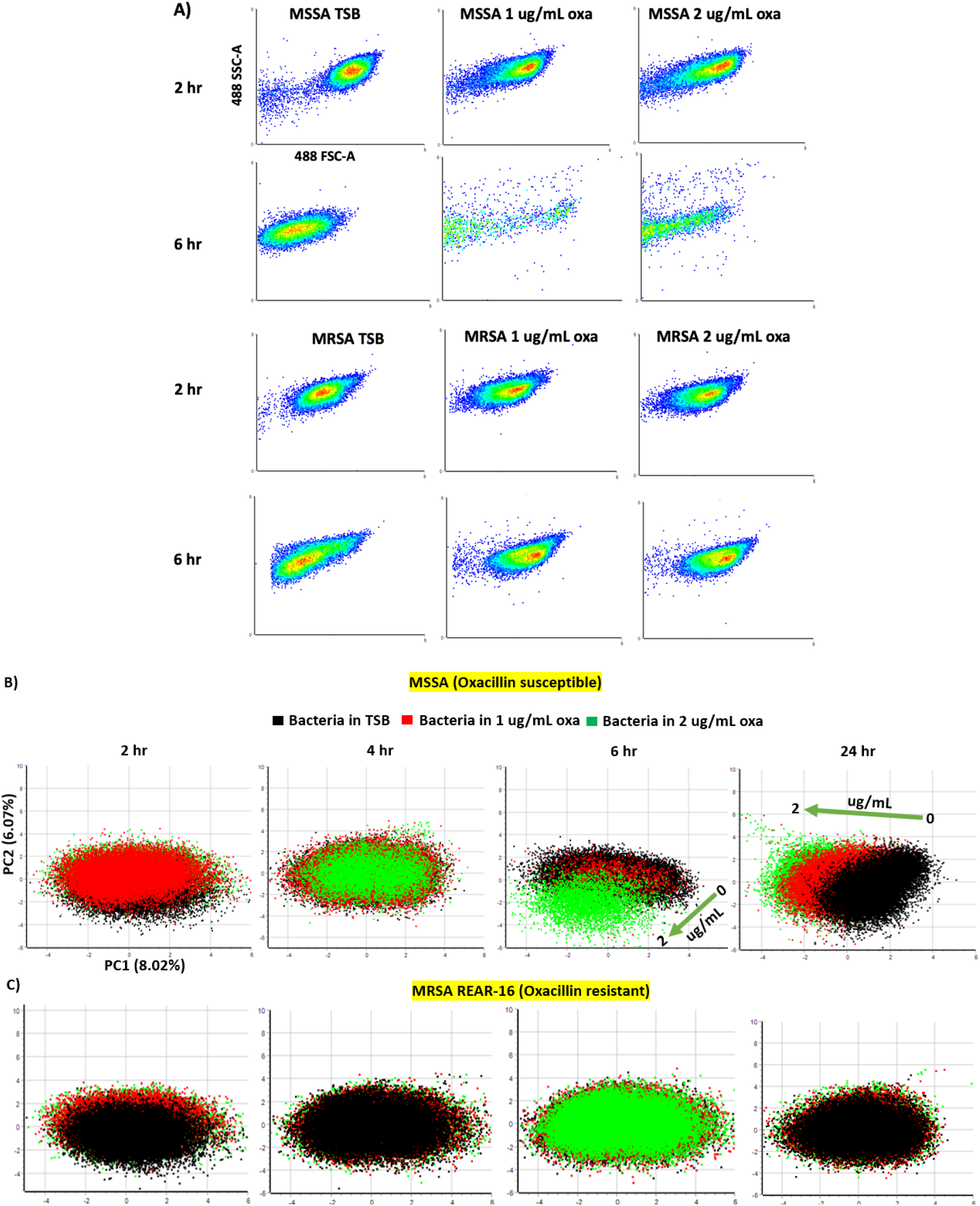
Scatter plots and PCA plots showing the effect of oxacillin stress on MSSA and MRSA autofluorescence.(A) Scatter plots showing the change in MSSA and MRSA cell density for non-stressed and stressed case (1 and 2 ug/mL Oxacillin). For MSSA stressing them with oxacillin shows more dying cell compared to the non-stressed case and more dead cells were observed with increasing concentration of oxacillin and incubation time. MRSA shows no significant cell death when stressed with oxacillin compared to the non-stressed case. (B) PCA plots of the non-stressed (black) and oxacillin stressed (1 ug/mL - red and 2 ug/mL - green) for MSSA over four different time points (2, 4, 6 and 24 h). It can be observed that at time points 2 and 4 h overlap in data clusters were observed for the stressed and the non-stressed case. After 6 and 24 h of incubation, a shift in the autofluorescence data points for oxacillin stressed samples (red and green) were observed. (C) PCA plots of oxacillin resistant MRSA showing no shift in the clustered data points for the stressed or the non-stressed case at four time points were observed. Oxacillin resistant and non-resistant Staph.aureus can be detected in 6 h by observing the difference in shift in autofluorescence signature.

Fig.5B and Fig.5C shows the PCA plots obtained from the autofluorescence spectra at selected wavelengths for both MSSA and MRSA at all the four incubation time points. It can be observed that for oxacillin susceptible MSSA, at incubation time points 2 and 4 h, there is a complete overlap in the data points for stressed (red and green) and non-stressed (black) cases. However, after incubating for 6 h, a clear shift in the data points were observed (green) for MSSA exposed to high concentration of oxacillin (2 ug/mL) compared to non-stressed case (black). This is also represented using a green arrow mark to represent the increase in oxacillin concentration at 6 and 24 h PCA plots. Similarly, after 24 h of incubation, shift in the oxacillin stressed MSSA (red and green) compared to the non-stressed MSSA (black) were observed. In the case of oxacillin resistant MRSA, no shift in the data points were observed for all the incubation times from 2 to 24 h. This shows that it is possible to differentiate between resistant and non-resistant *Staph*.*aureus* strain between 4 to 6 h by simply observing the shift in autofluorescence spectral signature (Intensity vs wavelength) or on the PCA plots for the antibiotic stressed bacteria.

### Comparing autofluorescence between different oxacillin resistant MRSA

Two more clinically isolated MRSA strains (GC-14 and CWND-19) were used, and experiments were repeated to measure the autofluorescence of stressed and non-stressed MRSA strains over four time points (2, 4, 6 and 24 h). Non-stressed MRSA strains (black) was used as positive controls. From Fig.5 we already established that the difference between MSSA and MRSA can be differentiated after 6 h of incubation with antibiotics. As an example, density plots obtained from analyzing autofluorescence signatures at selected wavelengths for two time points (2 and 6 h) for both the MRSA strains is shown in Fig.6. The data for time points 4 and 24 h is shown in supplementary Fig.S3. The density plots for both the strains showed no major cell death by stressing with oxacillin similar to MRSA REAR 16 strain in Fig.5A. This shows that all the three MRSA strains (REAR-16, CWND-19, and GC-14) are resistant to exposure to 1 and 2 ug/mL oxacillin after incubating for 6 h.

**Figure 6.**
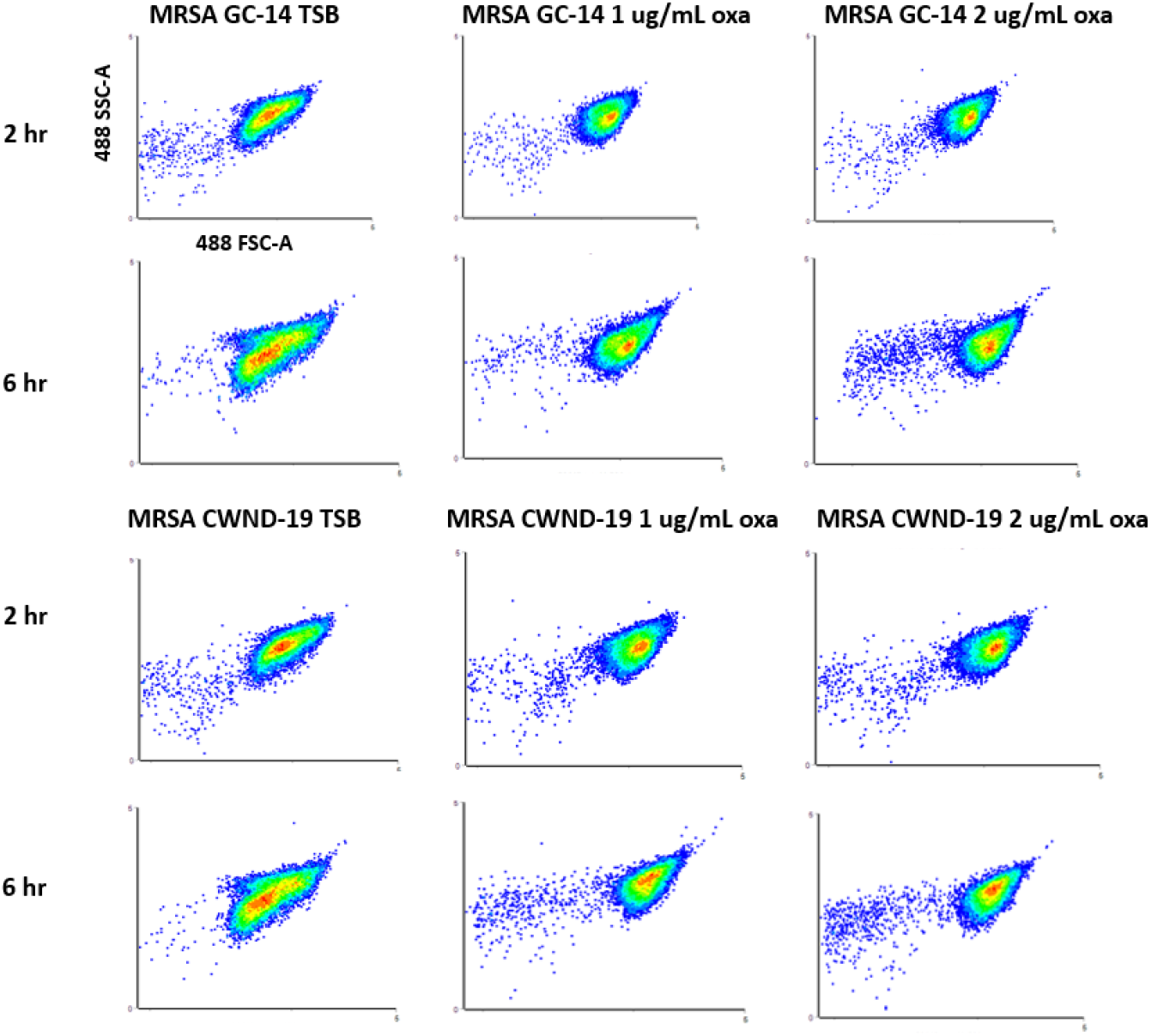
Scatter showing the effect of oxacillin on two clinically isolated MRSA strains. Scatter plots comparing two MRSA strains (MRSA GC-14 and MRSA CWND-19) for non-stressed and stressed case. No significant death of cells were observed on the scatter plot when exposed to1 and 2 ug/mL oxacillin at 2 and 6 h.

PCA analysis was performed on the autofluorescence spectral signature for both the MRSA strains and the PCA plots obtained are shown in Fig.7A and 7B. Similar to MRSA REAR-16 in Fig.5C, both GC-14 and CWND MRSA strains showed a completely overlapped data points in the PCA plots for non-stressed (black: TSB) and stressed (red : 1 ug/mL, green : 2 ug/mL) cases at incubation times 2 and 6 h. This observation reconfirms that, all the MRSA strains (REAR -16, GC-14 and CWND-19) which has developed resistant to oxacillin show no change in autofluorescence signature when exposed to oxacillin stress (1 and 2 ug/mL) and can be distinguished from the susceptible strains. The PCA plots of both the MRSA strains at 4 and 24 h is shown in supplementary Fig.S5.

**Figure 7.**
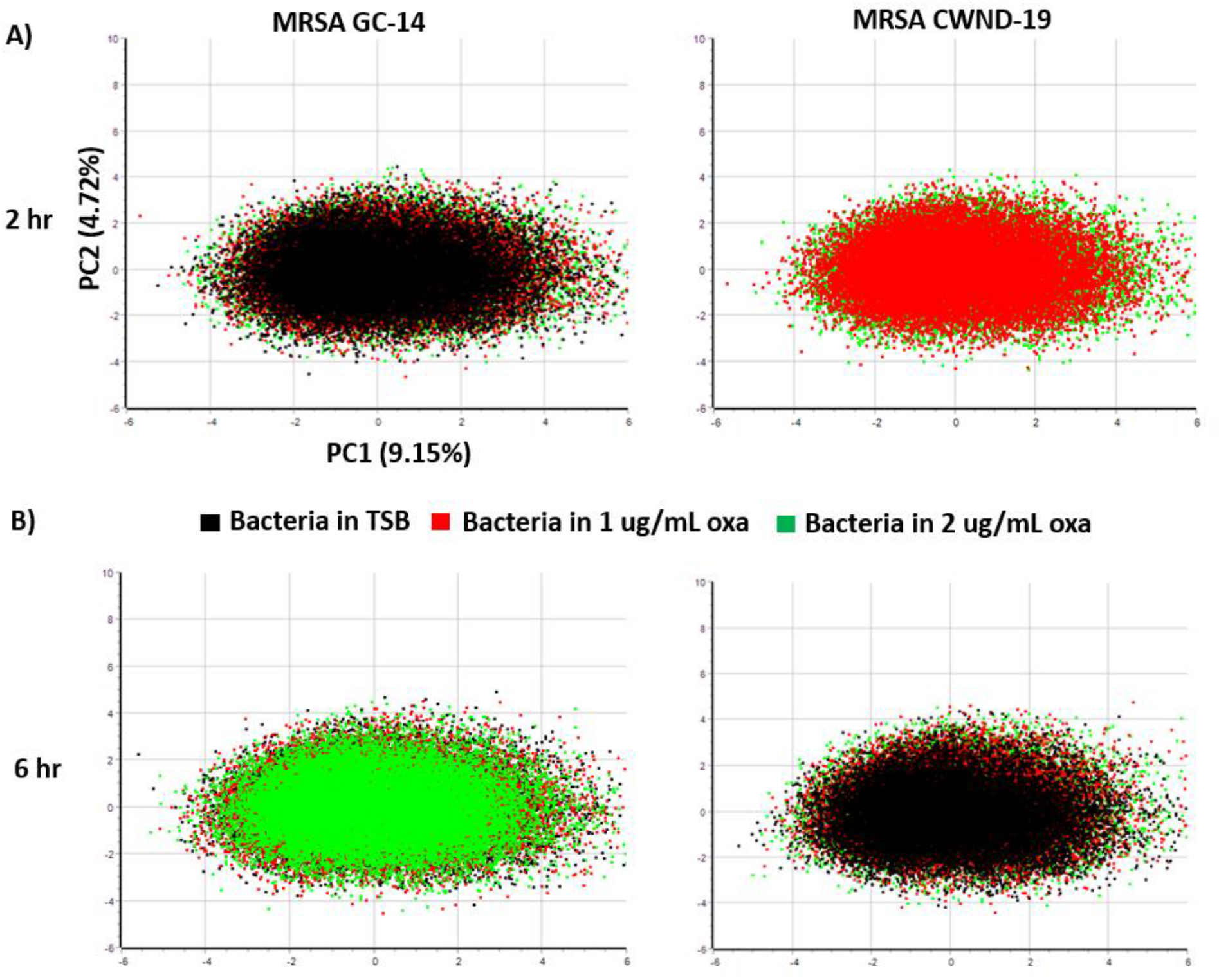
PCA plots showing the effect of oxacillin on two clinically isolated MRSA strains. (A) and (B) PCA plots for two MRSA strains after 2 and 6 h of incubation. A complete overlap of the data points for stressed and non-stressed cases were observed. There were no difference observed between oxacillin stressed and the non-stressed cases for both the MRSA strains.

## Discussion

Antibiotic resistance is a growing global concern which need urgent attention. Rapid diagnosis of pathogenic bacteria prevents overuse or misuse of antibiotics and promote specific treatment. Among them, MRSA is one of the most dangerous methicillin/oxacillin resistant strains causing a major threat to human life (Nandhini et al. 2022). A major problem in detecting MRSA is the presence of two sub-populations of antibiotic susceptible and resistant *staph aureus* in the same culture, termed as hetero-resistant (CDC 2019b). This hetero-resistant populations are observed in *staph aureus* that are resistant to oxacillin (CDC 2019b). In terms of treatment with antibiotics, presence of hetero-resistant can be highly challenging.

Standard phenotypic analysis methods employed to diagnose antibiotic resistant bacteria such as agar plating are very slow and it takes multiple subculturing steps (Vasala et al. 2020). This is mainly because sample containing suspected bacteria is initially plated for obtaining positive cultures. From the positive cultures, the colonies are grown again in the presence of antibiotics either by plating, E-test, or broth culture (CDC 2019b). As the result of these multiple culturing steps, the time taken to determine the antibiotic susceptibility (MIC values) can take up to several days depending on the starting sample type (Balouiri et al. 2016; Kowalska-Krochmal and Dudek-Wicher 2021).

Genomic analysis such as PCR are interesting alternatives for quick pathogen detection and identification. However, the sensitivity and specificity of PCR can be compromised depending on the starting sample type such as in beef samples where the innate property of the sample can hinder PCR amplification (Galhano et al. 2021). Moreover, PCR also needs pure samples such as from positive cultures for target specific amplification and PCR as a standalone method cannot differentiate between live and dead bacteria (Adzitey et al. 2013; Narayana Iyengar et al. 2021a; Narayana Iyengar et al. 2021b). The information about bacterial viability is crucial for infection treatment to avoid misuse or overuse of antibiotics. Other techniques such as MALDI-TOF can detect bacteria in a few minutes for bacterial identification by comparing the spectrum to the database (Patel 2015). Some of the main limitations of MALDI-TOF method are: it cannot differentiate between closely related bacterial species, needs positive bacterial cultures, and bacterial identification are limited to the reference spectra available in the database (Haider et al. 2023; Rychert 2019).

There are other upcoming technologies such as microfluidic platforms, single cell Raman spectroscopy and surface enhanced Raman spectroscopy (SERS) techniques that have gained significant interests as they are capable of performing single cell bacterial detection and identification (Li et al. 2012; Rebrosova et al. 2022). Despite the capabilities of performing sample preparation, detection, and identification of pathogens from varieties of samples, microfluidics technologies still suffer from the tradeoff between high throughput, efficiency, effect from inoculum, influence of sample matrix on assays (Iyengar et al. 2021; Needs SH 2020). Due to low throughput in microfluidic technologies, time to diagnose infection is high for applications such as blood stream infection and sepsis which demand high volumes of blood to be processed due to low concentration of bacteria (Liu et al. 2017; Narayana Iyengar et al. 2021b). On the other hand, single cell Raman spectroscopy provides very less signal from bacteria and thus SERS are mostly employed to obtain enhanced signals for bacterial identification (Rebrosova et al. 2022; Samek et al. 2021). However, signals from SERS can vary depending upon the type of starting sample, type of media used, interaction between sample and substrate, etc. which affects the sensitivity and specificity of detection (Rebrosova et al. 2022; Witkowska et al. 2020). In addition, all these techniques need a positive growth culture for further analysis which increases the total time needed for diagnosis (Rebrosova et al. 2022; Samek et al. 2021).

Flow cytometers and sorters are an interesting alternative diagnosis technique as they can detect and sort cells at a single cell level at high throughput and efficiency (Betz et al. 1984; Boye et al. 1983; Robinson 2021; Robinson 2022; Robinson et al. 2023). Bacterial analysis using conventional flow cytometers is usually performed by using staining methods which are either general stains or specific to bacterial species (e.g. antibody based, metabolism stains, protein binding stains etc.) (Davey and Kell 1996; Robinson et al. 2023; Shapiro 2003). However, staining methods are cumbersome as they involve multiple washing steps, stains can be expensive and availability of species-specific stains, especially specific to antibiotic resistant strains, are highly challenging (Drescher et al. 2021). In addition, the majority of the stains have broad emission signal which lead to signal overlap and compensation methods are employed to subtract the unwanted signal (Roederer 2001). If the peak emission signal overlaps completely then it is impossible to separate the signal and prevents multiplexing capabilities (Robinson et al. 2023; Robinson et al. 2005). One way to address this is to utilize natural autofluorescence signal from bacteria for detection. Autofluorescence is usually considered as a background noise and unmixed or subtracted from the main fluorescence. They are natural signals from bacteria which are emitted from metabolic proteins, enzyme co-factors (Flavin pathway) such as Flavin proteins, FAD and NADH etc. which are involved in bacterial metabolism (Kolenc and Quinn 2019; Skala et al. 2007).

For clinical applications, flow cytometers had to be kept inside a BSL-2 hood in order to handle pathogenic organisms, but many of the cytometers were very big to fit inside the hood and if they did, it made handling the samples uneasy (Robinson et al. 2023).Modernized flow cell sorters such as Bigfoot spectral cell sorters can address some of the limitations of the conventional flow cytometers. Bigfoot is equipped with 9 lasers and 64 detectors which not only has multiplexing capabilities but can also sort single cells at high throughput (70,000 cells/s) and efficiency for downstream analysis (ThermoFisher 2020). Bigfoot can handle a very high concentration of bacteria and can analyze at very high throughput at 70,000 cells per sec (ThermoFisher 2023). Bigfoot is qualified with biosafety cabinet similar to BSL-2 hood which can handle pathogenic organisms safely. As soon as the samples are introduced into the flow cytometers, the presence of bacteria can be identified directly by observing the bright field scatter plots or density plot (FSC vs SSC). By simply gating the bacterial population of cells, bacterial analysis can be performed on selected population.

In this report, we present an alternative approach for detection and identification of antibiotic resistant bacteria by measuring the natural spectral autofluorescence signatures using Bigfoot spectral flow cytometer. This is demonstrated by exposing the antibiotic susceptible and resistant bacteria with different concentration of oxacillin and by comparing their autofluorescence signatures over a few hours of incubation. As a proof of principle, we initially compared the spectral autofluorescence signature between non-stressed and gentamicin stressed bacteria (gentamicin susceptible *E*.*coli* and *Salmonella sp*.) We observed that gentamicin stressed *E*.*coli* and *Salmonella sp*. showed increase in autofluorescence intensity with increasing concentration of gentamicin at UV, blue and green excitation wavelengths and their corresponding emission signals recorded by 29 detectors. Increase in autofluorescence was observed in the intensity vs emission wavelength plot. After determining the specific wavelengths which showed increase autofluorescence in antibiotic stressed *E*.*coli* and *Salmonella* sp, the experiments were repeated using MSSA and human clinical isolate MRSA REAR-16 by stressing them with two different concentrations of oxacillin (1 and 2 ug/mL) and measured the autofluorescence signatures at four different time points (2, 4, 6 and 24 h). Oxacillin resistant MRSA showed no increase in autofluorescence at any time points. However, oxacillin susceptible MSSA showed an increase in autofluorescence at these specific wavelengths after 6 h of incubation for 1 and 2 ug/mL oxacillin concentration. This was observed by collecting intensity values from 20,000 cells per sample (about 1 minute). PCA analysis was performed on the raw spectral signature data and a clear shift in data clusters for the stressed bacteria (*E*.*coli*, SE for gentamicin and MSSA for oxacillin) were observed. We further confirmed this result by repeating the experiments with two more human clinical MRSA strains isolated (GC-14 and CWND-19) and no increase in autofluorescence were observed for any time points. An important thing to note here is that all these pathogenic organisms were analyzed in a normal laboratory settings where the Bigfoot spectral flow cytometer was operated due to the tight safety conditions within the instrument. In addition, by looking at the scatter plots (FSC vs SSC) we also observed that MSSA cells were dying when exposed to oxacillin with increase in incubation time, but it was not observed for MRSA. Increase in autofluorescence from bacteria is linked to an adoptive cellular response to external stress such as from antibiotics (Surre et al. 2018). Cells increase energy production in order to counter the damage from external stress which results in ATP (Adenosine Triphosphate) production. Monitoring the FAD and NAD production in eukaryotic cells are used to track the energy production (Bartolome and Abramov 2015; Heikal 2010; Surre et al. 2018). Flavin proteins are known to play an important role in DNA repair, energy metabolism, apoptosis, immune defense amongst others (Joosten and van Berkel 2007; Macheroux et al. 2011; Piano et al. 2017; Surre et al. 2018). As a result, the susceptible bacteria employs flavin based electron transport chain pathway and the concentration of FAD and NADH increases as a part of survival from antibiotic stress (Langer et al. 2013; Streker et al. 2005; Surre et al. 2018). However, resistant bacteria such as MRSA has the presence of resistant genes and can prevent the effect of certain concentration of antibiotic stress. Under stress, we also observed a slight increase in autofluorescence for MRSA with increase in oxacillin concentration, but it is not significant compared to non-resistant strain under oxacillin stress.

The main advantage of this approach is that the presence of antibiotic resistant strain in the sample was detected in 6 h compared to the conventional 72 h from agar plating techniques. The bacteria processed through the flow cytometers are still viable and can be used for further downstream analysis if needed such as PCR for genomic analysis. This prevents the need for agar plating and multiple culturing steps. The sample containing suspected bacteria can be exposed to the known antibiotic MIC values and their presence can be confirmed in few hours, making this approach a semi-quantitative method. In addition, no stains are needed for detection which can save time (prevents multiple washing steps) and money (cost of stains). As spectral signatures are utilized for AMR strain detection, more information can be obtained from the sample instead of specific peak emission signals in conventional flow cytometry. The sensitivity of detection is also not compromised in this method as there are no washing steps unlike staining techniques. The biosafety features in the Bigfoot allows safe handling of pathogenic bacteria in normal laboratory settings which was never possible in the field of flow cytometry before. Flow cytometers are already used in the clinical settings mostly for blood and cancer cell analysis and counting. This technique could be easily implemented in the clinical settings for AMR diagnosis from wide varieties of samples such as food, urine, blood among others.

## Author contribution

SNI, JPR, BR, and EB were responsible for conceptualizing and designing the research. SNI performed all the experiments and wrote the manuscript. BD and KR assisted with the usage and maintenance of the cell sorter. SNI, BD, and KR developed the working protocols for safely growing and maintaining the pathogenic organisms. BR and VP implemented developed the analysis software Cytospec 11.14 version. SNI, JPR, BR, and EB were responsible for reviewing and editing the manuscript. JPR, EB, and BR are responsible for funding and resource acquisition. All authors have read and approved the final version of the manuscript.

## Data availability

All data supporting the findings of this study are presented within the paper and the supplementary documents. The raw data, such as flow cytometry files (FCS), are available upon request.

## Ethical approval

This article does not entail any studies that involved human participants or animals.

