## Supplementary for "Measuring autofluorescence spectral signatures for detecting antibiotic-resistant bacteria using Thermofisher’s Bigfoot spectral flow cytometer"

### 1 Department of Basic Medical Sciences, Purdue University, West Lafayette, IN 47907, USA

### 2 Weldon School of Biomedical Engineering, Purdue University, West Lafayette, IN 47907, USA

### 3 School of Mechanical Engineering, Purdue University, West Lafayette, IN 47907, USA

### 4 Bindley Bioscience Center, Purdue University, West Lafayette, IN 47907, USA

**PCA plots using FCS express 7.0**

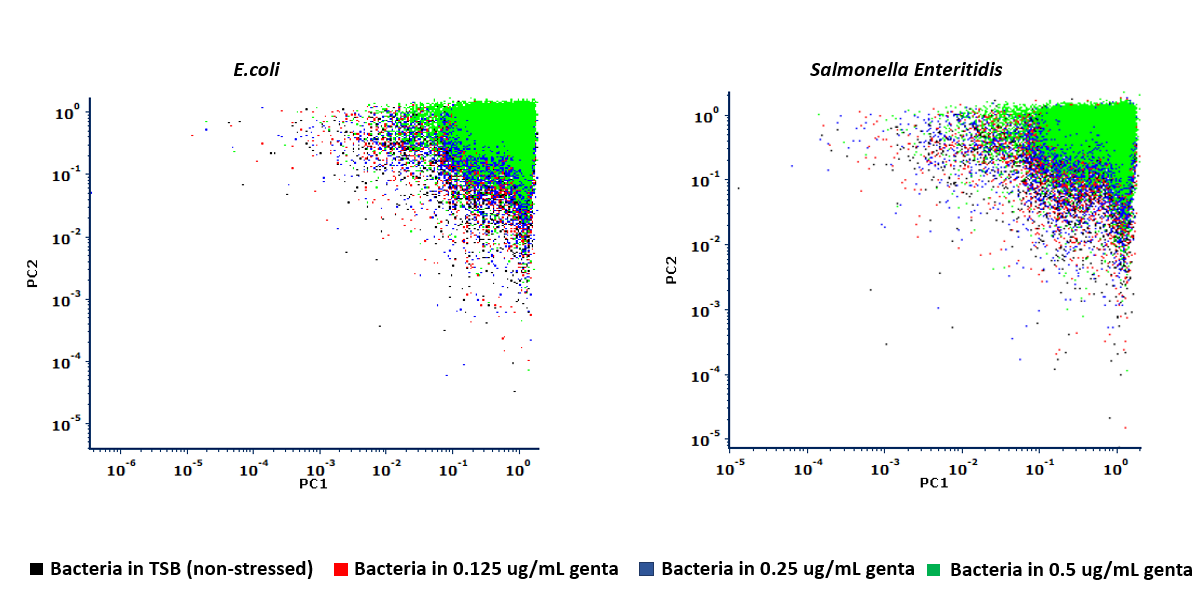
